# Genomic Transfer via Membrane Vesicle: A Strategy of Giant Phage phiKZ for Early Infection

**DOI:** 10.1101/2023.12.31.573766

**Authors:** Daria Antonova, Anna Nichiporenko, Mariia Sobinina, Innokentii E. Vishnyakov, Andrey Moiseenko, Inna Kurdyumova, Mikhail Khodorkovskii, Olga S. Sokolova, Maria V. Yakunina

**Affiliations:** Peter the Great St. Petersburg Polytechnic University, St. Petersburg, 195251, Russia; Institute of Cytology of the Russian Academy of Science, St. Petersburg, 194064, Russia; Faculty of Biology, Lomonosov Moscow State University, 1 Leninskie Gory, Bld. 12, 119234, Moscow, Russia; Faculty of Biology, Shenzhen MSU-BIT University, 1 International University Park Dr, Dayun New Town, Longgang District, Shenzhen, 518172, China

**Keywords:** phiKZ, phage nucleus, EPI vesicle, electron microscopy, electron tomography, fluorescent microscopy, membrane fluorescent dye, giant bacteriophage, *Chimaviridae*

## Abstract

During infection, the giant phiKZ phage forms a specialized structure at the center of the host cell called the phage nucleus. This structure is crucial for safeguarding viral DNA against bacterial nucleases and for segregating the transcriptional activities of late genes. Here, we describe a morphological entity, the early phage infection vesicle (EPI vesicle), which appears to be responsible for earlier gene segregation at the beginning of the infection process. Using cryo-electron microscopy, electron tomography, and fluorescence microscopy with membrane-specific dyes, we found that the EPI vesicle is enclosed in a lipid bilayer originating, apparently, from the inner membrane of the bacterial cell. Our investigations further disclose that the phiKZ EPI vesicle contains both viral DNA and viral RNA polymerase (vRNAP). We have observed that the EPI vesicle migrates from the cell pole to the center, displaying co-localization with ChmA, the primary protein of the phage nucleus. While phage DNA is transported into the phage nucleus after phage maturation, the EPI vesicle remains outside. We hypothesized that the EPI vesicle acts as a membrane transport agent, efficiently delivering phage DNA to the phage nucleus while protecting it from the nucleases of the bacterium.

## 1. Introduction

Giant bacteriophages stand out among bacterial viruses because of their huge genome (exceeding 200 kbp) and large particle size (approximately hundreds of nm in diameter) (1,2). Also, these bacteriophages exhibit unique infection development and structures. Virions of giant bacteriophages, especially the phiKZ-like phages, incorporate a cylindrical feature known as the inner body (1–3). This formation seems to play a role in packing the DNA within virions (2,4–6) and might have other functions, such as potentially facilitating DNA ejection or modulating the infection process (7,8). Moreover, giant bacteriophages such as 201phi2-1, phiKZ, and phiPA3 have evolved mechanisms to elude DNA-targeting bacterial defenses (9,10). Namely, they form a proteinaceous shell at the cell center during infection, termed the phage nucleus, which encapsulates their viral DNA (9,11). Inside the phage nucleus, the DNA replication (12) and transcription of middle and late phage genes (13) occur. The main protein of the shell was determined and named ChmA (short form from Chimallin) (14), which identifies the new *Chimaviridae* phage group encompassing phiKZ-like phages, Goslar and SPN3US phages (15). The architecture of the phage nucleus shell reveals pores that permit the passage of smaller molecules like RNA and DNA (14,16,17). Still, the transit of larger proteins associated with DNA replication, transcription, and recombination (18), as well as proteins fused with a GFP mutant and YFP (19,20), remains an enigma. To place this structure centrally within the cell, these bacteriophages produce tubulin-like proteins, PhuZ, that assemble into a spindle structure (9,21). Also, the spindle guides the viral capsids from the peripheries of the cell to the phage nucleus surface for DNA encapsulation (22).

Interestingly, the phiKZ phage nucleus consistently safeguards its viral DNA from restriction-modification and CRISPR-Cas DNA-targeting systems, even those atypical for *P. aeruginosa* hosts (10). This suggests a universal resistance strategy by phiKZ against diverse bacterial defense systems. However, the phage nucleus forms during the mid-infection stage, leaving the protection of phage DNA during the early infection phase a topic for study. Earlier we identified round compartments (RC) forming at the beginning of phiKZ infection (11). Also, similar compartments were found for some other giant bacteriophages such as SPN3US, 201phi2-1 and Goslar (12,14,23). Recently, these structures were shown to be surrounded by a lipid bilayer in the case of bacteriophages 201phi2-1 and Goslar (12,14). These have been termed early phage infection (EPI) vesicles. In this research, we confirm the membranous nature of the phiKZ EPI vesicle boundary and demonstrate that the phiKZ EPI vesicle (or RC as we previously named it) contains phage DNA in the early stage of infection. Also, using live fluorescent microscopy, we revealed that the EPI vesicle took part in transferring phage DNA to the middle of the cell with ChmA. We propose that the phiKZ EPI vesicle acts as a safeguard for the viral DNA against bacterial defenses in the initial infection phase, facilitates DNA migration to the cell’s center, and also plays a role in transferring DNA to the phage nucleus.

## 2. Materials and Methods

### 2.1 Bacteriophage, Bacterial Strain and Growth Conditions

*P. aeruginosa* PAO1 culture was grown in LB medium at 37 °C with aeration at 200 rpm. Cells were diluted 1:100 in fresh LB medium and grown to an optical density at 600 nm (OD600) of 0.6. At this point, the culture aliquots were used either for microscopic experiments or for obtaining phage yield.

*P. aeruginosa* PAO1 cells were transformed with a plasmid expressing fusion protein genes as described in our previous study (24) and selected on LB with 1,5% agar plates containing carbenicillin (100 ug/ml). A single transformed colony was grown overnight at 37 °C in liquid LB medium supplemented with 100 μg/ml carbenicillin. The culture was diluted 1:100 in fresh LB medium containing carbenicillin. PAO1 cells carrying plasmids were grown at 37 °C until OD600 reached 0.2 and induced with arabinose (final concentrations of 0.2% for pHERDmChGp180 and 0.02% for pHERDmChGp54). Growth continued at 30 °C until OD600 reached 0.6. At this point, culture aliquots were used for fluorescent microscopy experiments or for phage infection to obtain phage progeny containing fusion fluorescent proteins.

High-titer phiKZ preparations were prepared from lysed infected PAO1 cultures and purified by centrifugation at 10000g for 30 min. Phage particles were purified and concentrated from the supernatant by PEG-8000 as described in the article (25).

### 2.2 Cloning

Plasmid pHERDmChgp54 is the derivative of the shuttle vector pHERD20T. Firstly, mCherry fluorescent protein gene was inserted into pHERD20T vector according to the manufacturing protocol of NEBuilder® HiFi DNA Assembly kit (New England Biolabs, UK). The resulting plasmid was referred to as pHERDmCh. Then Gp54 gene (encoded Chimallin, ChmA) was PCR amplified from phiKZ genomic DNA and cloned into pHERDmCh by EcoRI/HindIII and into pET28 by NdeI/HindIII restriction sites. The resulting plasmids were named pHERDmChGp54 and p28HisGp54. Plasmid pHERDmChGp180 was generated previously (13).

### 3.1 Fluorescent microscopy

The *P. aeruginosa* PAO1 cells at OD600 of 0.6 were infected by bacteriophage phiKZ with multiplicity of infection (MOI) = 10. Lipophilic dye FM4-64 at a final concentration of 1.6 μM was added to cells in 5 min post-infection. To stain the inner cell membrane of *P. aeruginosa* PAO1 by FM4-64 dye, 1 mM or 10 mM EDTA was used as an outer membrane permeabilizer. In the case of live cell imaging, Mitotracker Green FM dye (Thermofisher, USA) was added to *P. aeruginosa* PAO1 cells (1:100 dilution of overnight culture) directly. Prestained PAO1 cells at OD600 0.6 were centrifuged at 5000 g for 5 min, the supernatant was removed and the pellet was resuspended in the same volume of liquid LB medium. These PAO1 cells were infected by native phiKZ bacteriophage or phiKZ bacteriophage including mCherry-gp180 protein in virions (MOI=10). PAO1 cells carrying pHERDmChGp54 were prepared similarly as described above.

Infected cells were placed on an agarose pad (0.25 LB medium, 1 % agarose) supplemented by DAPI DNA dye at 1 μM concentration. When cells were completely absorbed, the agarose pad was sealed by a cover glass and VALAP (lanolin, paraffin and petrolatum 1:1:1). The slide was mounted in a Nikon Eclipse Ti-E inverted microscope equipped with a custom incubation system. Filter sets TxRed-4040C, YFP-2427B and DAPI-50LP-A (Semrock) were used for red, green and DAPI fluorescence detection, respectively. Live cell imaging was performed using Micro-Manager with a custom script at 30 °C. Images were taken every 10 min for 180 min. Image analyses were performed using ImageJ.

### 4.1 Antibodies production

Rosetta(DE3) *E. coli* cells were transformed with pET28aHisGp54. Expression was induced by the addition of 1 mM IPTG to cultures grown to OD600 = 0.5-0.7 and further growth at 22°C for 3 hr. 2 g of wet biomass was disrupted by sonication in 20 ml of buffer A (420mM Potassium Phosphate Buffer pH=8,0; 10% glycerol; 500mM NaCl; 1mM DTT; 50mM L-Glu; 50 mML-Arg; 10 mM imidazole) followed by centrifugation at 11000g for 30 min. Clarified lysate was loaded onto a HisTrap HP 5 ml (GE Healthcare Life Sciences, USA) column equilibrated with buffer A and washed with the same buffer. The recombinant proteins were eluted with buffer B (buffer A containing 250 mM imidazole). The eluted fractions were concentrated on Amicon Ultra-4 Centrifugal Filter Unit with Ultracel-10 membrane (EMD Millipore, Merck, USA) to 3 mg/ml was used for rabbit immunization.

As a result, we obtained a 10 ml antiserum sample. The Anti-Gp54 antibodies (ABs) were purified using 1 ml of CnBr-activated Sepharose (GE LifeScience, USA) conjugated with purified His-Gp54 according to manufacturer recommendations. The purified ABs were eluted with 0.2 M glycine pH 2.8, into a microtube with the addition of 0.1 M Tris pH 8.8 and 0.3 M KCl. Then, the fractions containing ABs were combined and the buffer was changed to PBS buffer using Amicon Ultra-4 30 kDa (Millipore EMD, USA). AB concentration was determined using Bradford dye and glycerol was added up to 50% for storage at -20C.

### 5.1 Immuno-labeling assay

Samples of non-infected and phiKZ-infected cells immediately after phage addition (0-2 min), on 15 and 30 min of infection were chemically fixed using a mixture of glutaraldehyde (0.1%) and formaldehyde (2%) for 30 min at room temperature. The cells were collected by centrifugation at 5000g at 4°C. Then, cell pellets were washed twice with sterile PBS and subjected to LR White (Polyscience, Inc., USA) embedding, according to the manufacturer’s recommendations (Sigma-Aldrich, United States). Briefly, samples were dehydrated with increasing ethanol concentrations (70%, 95%) and then impregnated with the resin-alcohol mix in ratios 1:3 (2 hr, RT), 1:1 (overnight, 4 °C), 3:1 (2 hr, RT), and in the pure resin during 2 hr (RT). The resin-impregnated precipitates were placed in gelatin capsules (size 2, Electron Microscopy Sciences, United States), filled with fresh resin, and left to polymerize at 54 °C (2 days). Ultrathin sections were prepared using the LKB-III ultramicrotome (LKB, Sweden) and applied to nickel grids (400 mesh, Ted Pella, United States) covered with a collodion film (Sigma-Aldrich, United States). For immune electron microscopy, sections were blocked with 1% aqueous solution (or 1% solution in PBS) of bovine serum albumin, treated with the corresponding primary antibodies against the protein of interest ChmA (Gp54) (rabbit polyclonal antibodies) at least 4 hr, but no more than 8 hr for optimal binding of antibodies to the sample. Then, the grids with samples were washed 4 times in 1×PBS and incubated for 1 hr with secondary antibodies (goat anti-rabbit antibodies conjugated to colloidal gold particles with a diameter of about 15 nm, Aurion, Netherlands). The washing was repeated similarly to the previous step, and then the sections were stained with Uranyl Acetate Alternative (Gadolinium Triacetate based stain, Ted Pella, United States) for 10 min and viewed in a Libra 120 electron microscope (Carl Zeiss, Germany) at magnifications of 10000-16000×.

### 6.1 Transmission Electron Microscopy (TEM) and Electron Tomography

Overnight culture *P. aeruginosa* PAO1 was diluted 1:100 in fresh LB medium and grown at 37 °C to OD600 of 0.6. The bacterial cells were infected by phiKZ bacteriophage with MOI = 10. After 5 and 20 min of the infection, cells were collected and placed on ice to lower infection development before grid preparation.

The transmission electron microscopy and electron tomography studies were carried out with JEOL JEM-2100 200 keV electron microscope equipped with LaB6 electron source. The images were obtained either with the Gatan Ultrascan 1000XP CCD camera or with the Gatan Ultrascan 1000FTXP CCD camera mounted after the Gatan Imaging Filter (GIF Quantum). The energy filter was used in the ZLP mode to filter out inelastically scattered electrons during plastic section tomography acquisition.

Electron tomography data were collected with SerialEM software with ±60° range and 1° step. The areas of interest were baked with an electron dose higher than 1000 e/Å^2^ prior to the data collection to reduce the section shrinking during imaging. The tomograms were reconstructed with IMOD software (26) following the standard procedure.

### 7.1 Cryo-EM

The *P. aeruginosa* cells were infected with phiKZ immediately before grid preparation to keep the infection timings. 3.5 μl of the infected cells suspension was applied to the carbon side of glow-discharged lacey carbon films, and an additional 0.5 μl were applied to the back side of the grid. The grid was blotted for 12 sec from the back side of the grid using the Leica EM GP2 plunger and immediately vitrified in liquid ethane. Images of the whole vitrified cells were acquired with JEOL JEM-2100 200 keV electron microscope using Direct Electron DE-20 detector. The images were taken with -20 μm defocus at low magnification yielding a 7.2 Å pixel size and total electron dose of less than 10 e/Å^2^ to obtain a high signal-to-noise ratio, while keeping the sample intact.

## 3. Results

### 3.1 The phiKZ EPI vesicle is surrounded by a lipid bilayer

Electron microscopy tomography was carried out on the plastic sections of embedded *P. aeruginosa* cells infected with the phiKZ phage to visualize the structural peculiarities of round compartments (Fig. 1 A,C). Most of the EPI vesicles contain dense branching structures that occupy the majority of the space inside, at 5 min after phiKZ infection. From the tomogram cross-sections, it is clear that each vesicle is bordered with a bilayer membrane of around 5 nm thick, which corresponds to the lipid bilayer thickness (Fig. 1 B,D). The thin sections were stained with osmium tetroxide, giving membranes enough contrast to resolve in the plastic section tomography reconstructions.

**Figure 1.**
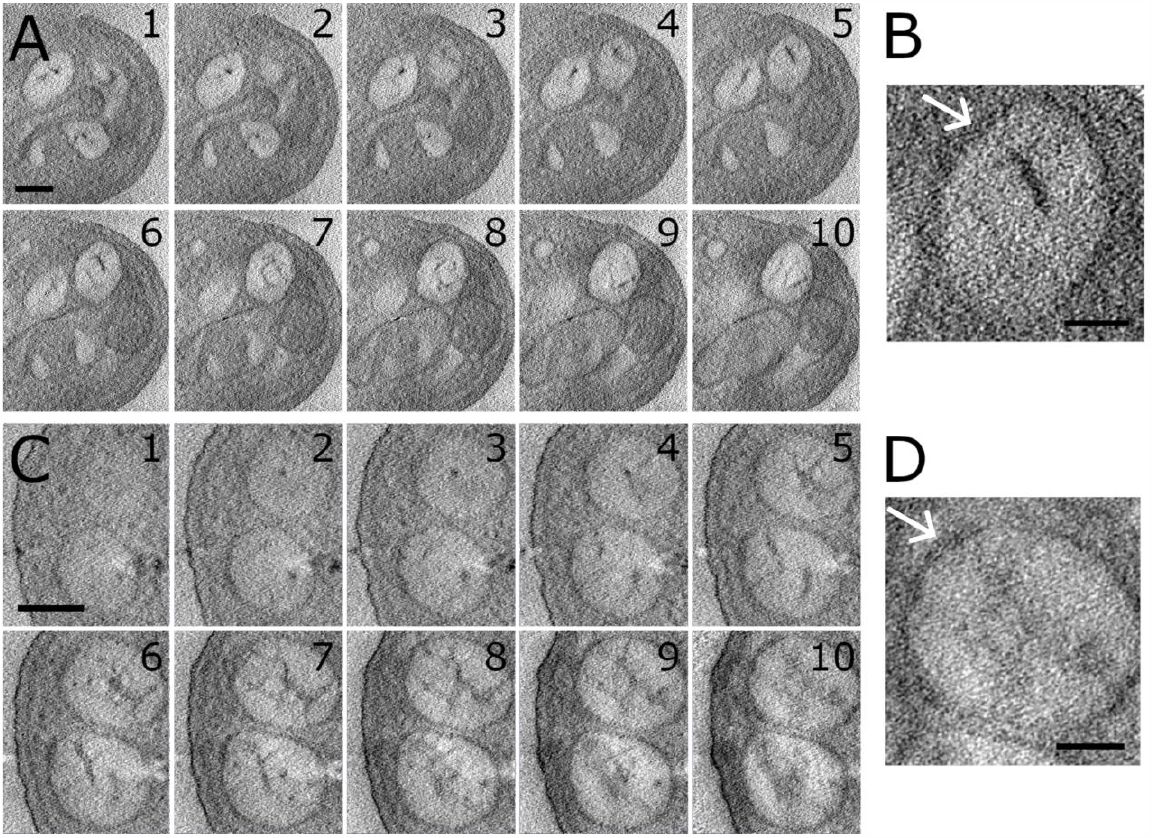
Z-slices through electron tomography reconstruction of the EPI vesicles in *P*.*aeruginosa* plastic sections at different times after infection: (A) - 2 min, (C) - 5 min. The branching electron dense inner structure is visible in each vesicle. Scale bar - 100 nm.; Zoomed views display the bilayer membrane bordering the EPI vesicles (white arrow): (B) - 2 min, (D) - 5 min. Scale bars - 50 nm.

Cryo-electron microscopy of the whole vitrified *P. aeruginosa* cells allowed to image of the EPI vesicles at different stages of phiKZ phage infection. The cells were vitrified after 5 and 20 min of infection. The EPI vesicles were found at both time points, close to the phiKZ phages infecting the cells (Fig. 2). In the 5 min samples the EPI vesicles are located near the inner cell membrane; in the 20 min sample, the EPI vesicles moved closer to the center of the cell. This observation correlates well with our previous data (11).

**Figure 2.**
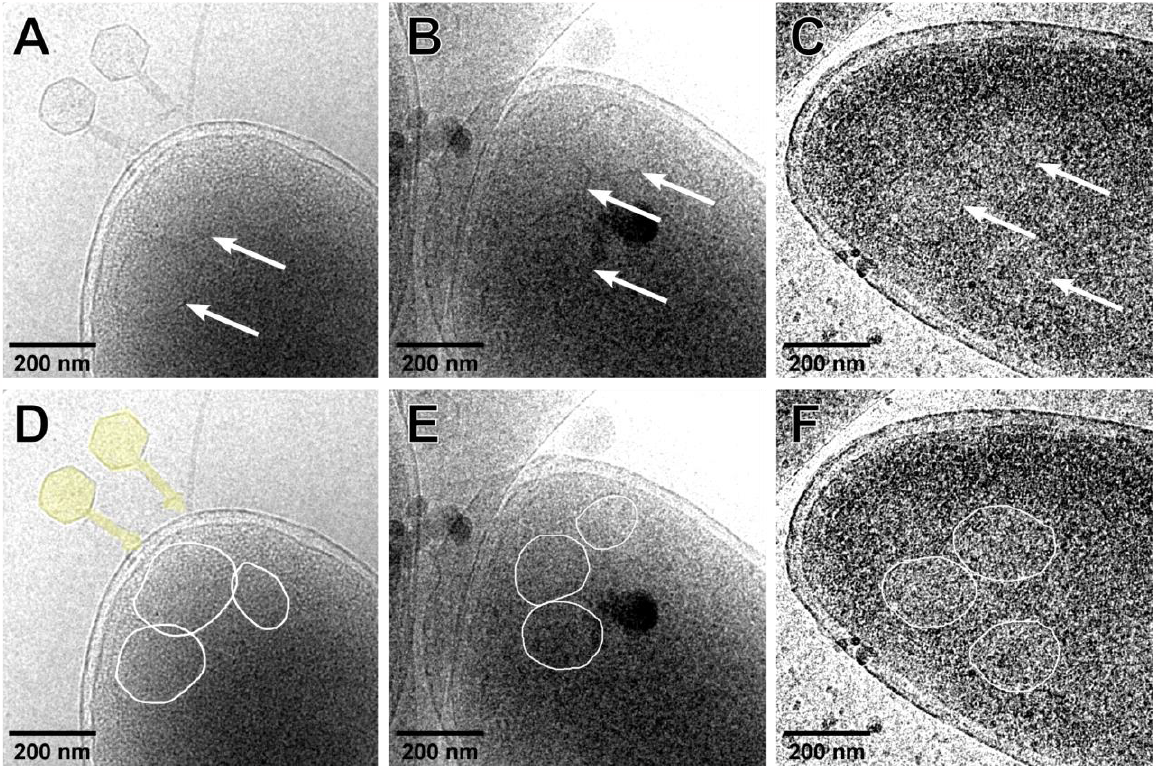
Representative cryo-EM images of EPI vesicles in *P. aeruginosa* cells infected with phiKZ bacteriophage and vitrified after 5 min (A,B) and 20 min (C) of infection. The original image contrast was changed for better display. B, E,F - corresponding images with EPI vesicles encircled with white lines for clarity; phage particles are highlighted yellow.

To further prove that phiKZ EPI vesicles are surrounded by the lipid bilayer, we employed lipophilic fluorescent dye FM4-64. Due to its amphiphilic character, it is incorporated into the outer membrane of gram-negative bacteria and is commonly used for membrane staining *in vivo* (27,28). We treated *P. aeruginosa* PAO1 cells with FM4-64, infected the cells by phiKZ, and placed them on an agarose pad with DAPI dye. As a result, EPI vesicles were not stained by FM4-64 in infected *P. aeruginosa* cells (Fig. S1). Based on these results, we suggest that the outer membrane does not take part in EPI vesicle formation. Next, we treated *P. aeruginosa* cells by FM4-64 in the presence of EDTA to permeabilize the outer membrane and allow the staining of the inner bacterial membrane (29). After phage addition, DAPI stained puncta corresponding to phage DNA (11) in phiKZ-infected cells were colocalized with puncta stained with FM4-64 dye. (Fig. 3). So, we speculate that the boundaries of EPI vesicles are formed by the inner membrane.

**Figure 3.**
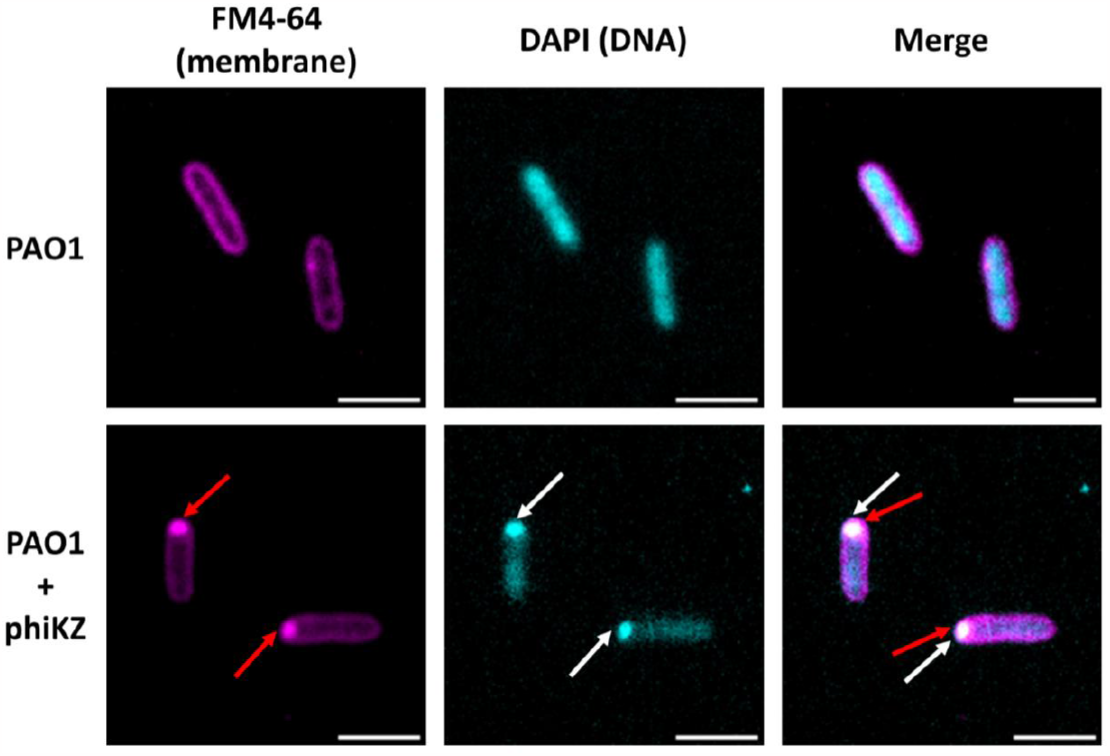
Fluorescence microscopy images of EPI vesicles stained by FM4-64 in the presence of 10 mM EDTA, colocalized with the DAPI-stained phiKZ DNA. Uninfected *P. aeruginosa* PAO1 cells as control are present in the first row, and phiKZ-infected PAO1 cells are in the second row. The EPI vesicles are indicated by red arrows, phiKZ DNAs are indicated by white arrows. Scale bar is 2 um.

Notably, the addition of EDTA prevented further infection development in the cell. It is likely due to EDTA binding divalent cations (30), which are required as cofactors for phage transcription enzymes (31), and/or this EDTA concentration renders an antibacterial effect and leads to cell death (32).

### 3.2 In the course of infection, the EPI vesicle moves to the cell center along with DNA

To localize EPI vesicles in live cells during phiKZ infection, Mitotracker Green dye was used to stain PAO1 cells. This dye is able to penetrate the outer membrane of the bacterial cells and stain the inner cell membrane (33,34) by covalently binding free sulfhydryl groups in proteins (35). PAO1 cells incubated with Mitotracker Green were subsequently infected by phiKZ and placed on an agarose pad containing DAPI dye. As a result, both DNA and inner membrane structures were observed in cells during infection. As can be seen in Fig. 4, the EPI vesicle punctum in Mitotracker Green channel colocalizes with DNA punctum near the cell pole as the infection begins. At 20 min post-infection, the EPI vesicle was found to move along with DNA punctum toward the cell center. Interestingly, after 30 min of infection, while the intensity of the DAPI dye signal increases in the center of the cell, implying the viral DNA replication in the phage nucleus, the EPI vesicle punctum remains close to the cell membrane and does not intersect with the DAPI signal. We suggested the EPI vesicle function is to be the DNA carrier to maintain phage DNA at the beginning of infection, before the maturation of the phage nucleus.

**Figure 4.**
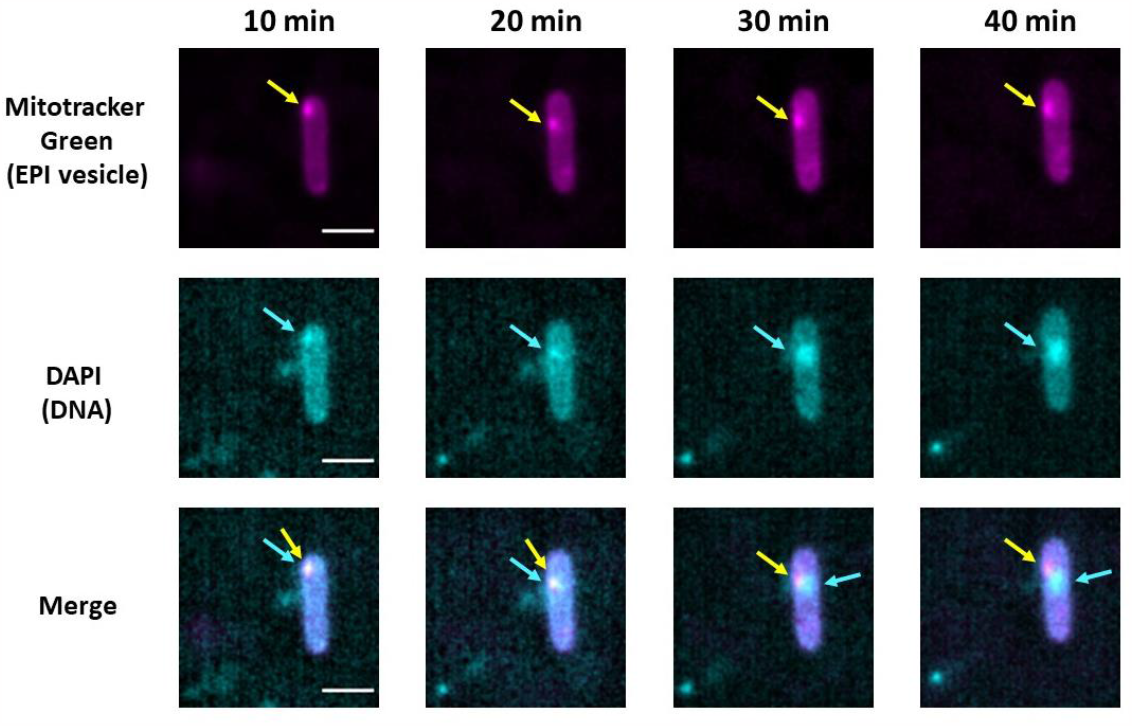
Fluorescence microscopy of the PAO1 cell infected by phiKZ phage and stained by the Mitotracker Green and DAPI dyes. The EPI vesicle is indicated by yellow arrows, viral DNA is indicated by blue arrows. The channel with Mitotracker Green stain (EPI vesicle) is presented in the top row, DNA channel is presented in the middle row, and the composite of the DNA and Mitotracker Green channels is presented in the bottom row. Min post-infection are indicated above the respective column. Scale bar is 2 um.

### 3.3 EPI vesicle lifecycle and ChmA location during phage nucleus formation

The main protein of the phage nucleus is ChmA, it is synthesized from the early phage gene soon after the start of infection (36). Earlier it was shown that ChmA-puncta (Gp54 protein, Chimallin) is migrating from the cell pole to the center along with the DNA-puncta (9,10). Our experiments reveal that originally the DNA is surrounded by an inner cell membrane (Fig. 3). To explore ChmA’s location in relation to the EPI vesicle, we used the cells, which synthesized the recombinant mCherry-ChmA fusion protein, and infected them with phiKZ. At 10 min post-infection, the EPI vesicle puncta stained by Mitotracker Green and mCherry-ChmA were localized close to each other. Yet at 20 min post-infection, the EPI vesicle and the formed phage nucleus seem to be separated. Notably, the EPI vesicle was not destroyed and later was found near the ChmA shell of the phage nucleus (Fig. 5).

**Figure 5.**
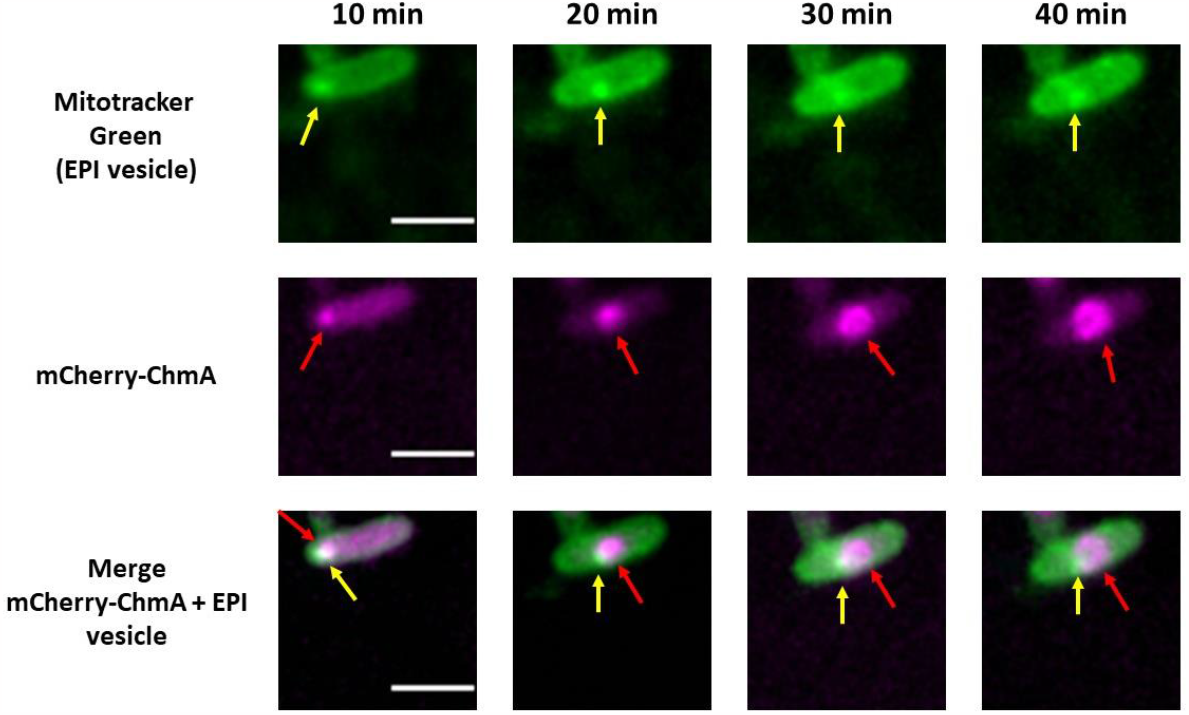
Fluorescence microscopy of the infected PAO1 cell synthesizing the mCherry-ChmA fusion protein. The EPI vesicle is indicated by yellow arrows, mCherry-ChmA accumulation puncta are indicated by red arrows. The EPI vesicle in the Mitotracker Green channel is presented in the top row, mCherry-ChmA is presented in the second row, and the the combination of the mCherry-ChmA and Mitotracker Green (EPI vesicle) channels is presented in the bottom row. Min post-infection are indicated above the respective column. Scale bar is 2 um.

We employed TEM to visualize the phage ejection and simultaneous EPI vesicle formation (Fig. 6, 0-2 min). To analyze the position of ChmA inside the cell, we used rabbit polyclonal antibodies (ABs) against this protein labeled with nanogold particles (Fig. S2). Immediately after phiKZ phage addition to the cell culture, the black dots, which indicate the ChmA position, were not detected (Fig. 6, 0-2 min). At the 15^th^ min, the ChmA-associated signal was revealed inside the cells, some of the dots were located near the EPI vesicles (Fig. 6, 15 min). At the 30^th^ min of infection, ChmA-associated signals were mostly located along the outer border of the mature phage nucleus, formed by ChmA (9,14). Simultaneously, we observed the EPI vesicle in close proximity to the phage nucleus, which corresponds with the above mentioned fluorescent data and our previous observations (11).

**Figure 6.**
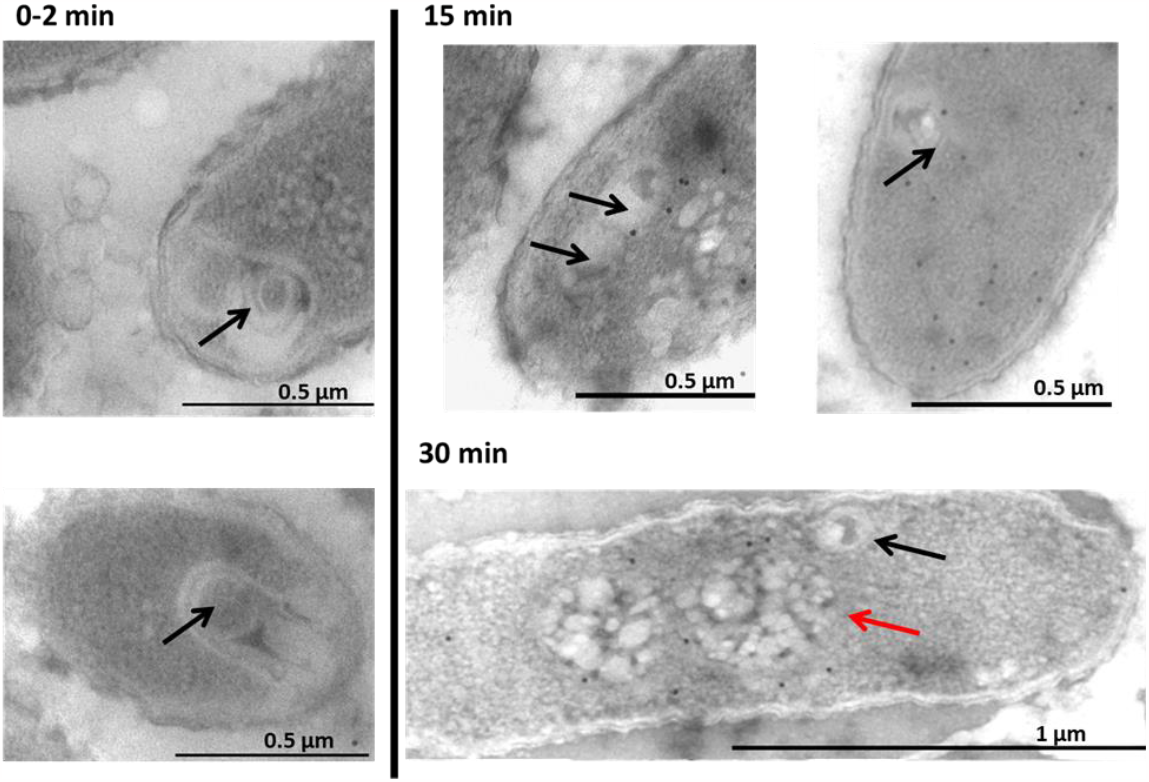
Localization of ChmA protein (Gp54 for phiKZ) in P. aeruginosa infected cells at 0-2, 15 and 30 min infection time points, determined by the immuno-gold labeling assay. Black arrows indicate the EPI vesicle position. Black dots correspond to gold particles associated with the ChmA. The red arrows indicate the phage nucleus. The black lines in the lower right corner represent scale bars. More examples are provided in the Supplementary file (Fig. S2).

### 3.4 PhiKZ vRNAP protein Gp180 is incorporated into the EPI vesicle

Gp180 protein is the subunit of virion RNAP (vRNAP), which is injected into the bacterial cell together with the phage DNA and promotes transcription of the phage phiKZ early genes (13,36). We decided to define the relationship between injected vRNAP and EPI vesicles. For that, phage particles containing labeled mCherry-gp180 vRNAP subunit were used to infect the PAO1 cells stained by Mitotracker Green. Fig. 7 shows that 10 min after infection, mCherry-Gp180 clearly colocalizes with the EPI vesicle below the cell membrane. When the EPI vesicle is displaced to the cell center (20 min), it is followed by mCherry-Gp180. At the 30 min of infection, EPI vesicles with vRNAP remain together outside the mature phage nucleus with replicating DNA. After 40 min of infection, fluorescence intensity of mCh-Gp180 substantially reduced. This can be caused by photobleaching or by the mCherry-Gp180 diffusion over the cell due to final disruption of the EPI vesicle.

**Figure 7.**
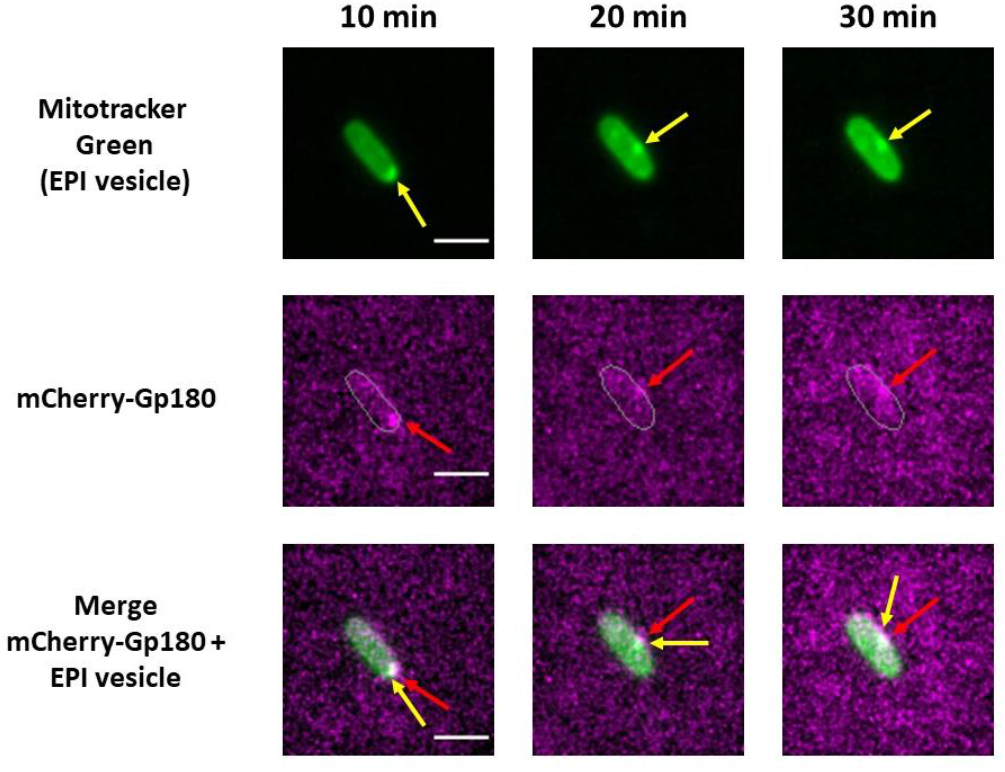
Fluorescence microscopy of the PAO1 cells infected by phage particles contains mCherry-Gp180. The EPI vesicle is indicated by yellow arrows, mCherry-Gp180 is indicated by red arrows. Mitotracker Green stain (the EPI vesicle) is presented on the top row, mCherry-Gp180 is presented on the second row, and the composite of the mCherry-Gp180 and Mitotracker Green channels is presented on the bottom row. Min post-infection are indicated above the respective column. Scale bar is 2 um.

## 4. Discussion

This study shows that the round compartments (RC) demonstrated by us earlier (11) have a boundary formed by a lipid bilayer and a dense branching internal structure. Apparently, this structure consists of packed viral DNA with some virion proteins such as vRNAP. Given the resemblance of RC to EPI vesicles observed in Goslar-infected cells (11,12), we have adopted the term ‘EPI vesicles’ to describe these compartments thereafter.

We concluded that the EPI vesicle is formed by the inner cell membrane invagination, based on the ability of FM4-64 dye, which *in vitro* stains the outer cell membrane of the intact gram-negative bacteria (34), to stain EPI vesicles after increasing its membrane permeability by EDTA. This is also supported by the fact that transport of EPI vesicles inside infected cells was detected using the Mitotracker Green dye, which can stain the inner cell membrane (33,34). In the course of infection, the EPI vesicles moved from the cell pole to the cell center and were co-localized with DAPI-puncta corresponding to phage DNA and with ChmA (Gp54 for phiKZ), fused to fluorescent mCherry protein. We showed that when the ChmA-shelled nucleus formed and the DAPI signal increased inside, which reflected phage DNA replication, the EPI vesicle was located outside the phage nucleus border, encapsulating the vRNAP. Interestingly, not only vRNAP, but other virion proteins: Gp90, Gp93, Gp95, and Gp97 enter the cell together with viral DNA (8). Moreover, all these proteins move inside an infected bacterial cell together with DAPI-puncta similar to EPI vesicle movement (7,8). Apparently, all these proteins are encapsulated inside the EPI vesicle.

Thus, our findings lead us to propose a three-stage model of early phiKZ infection (Fig. 8). In the initial stage, phiKZ injects its viral DNA and some virion proteins into the host cell, and a lipid bilayer, presumably derived from the inner cell membrane, encapsulates them, forming an EPI vesicle. This encapsulation immediately safeguards the viral components against bacterial defense mechanisms, facilitating the transcription of phage DNA by vRNAP. Inside the EPI vesicle, the early gene gp54 (corresponds to ChmA) is transcribed, and its mRNA transports to the cytoplasm for translation. In the second stage, ChmA appears to accumulate in the bacterial cytoplasm near the EPI vesicle and migrates with the EPI vesicle toward the cell center. In the third stage, ChmA in the cell center forms the phage nucleus. Given the evidence that DNA replication does not occur within the EPI vesicle (12), the phage DNA transfers into the phage nucleus for replication during this stage. However, the EPI vesicle remains near the nucleus, containing injected proteins. After the third stage, the development of phiKZ infection goes to the second part of the infection, which is based on the DNA transcription by nvRNAP inside the phage nucleus and maturation of phage progeny (13, 22, 36, 37).

**Figure 8.**
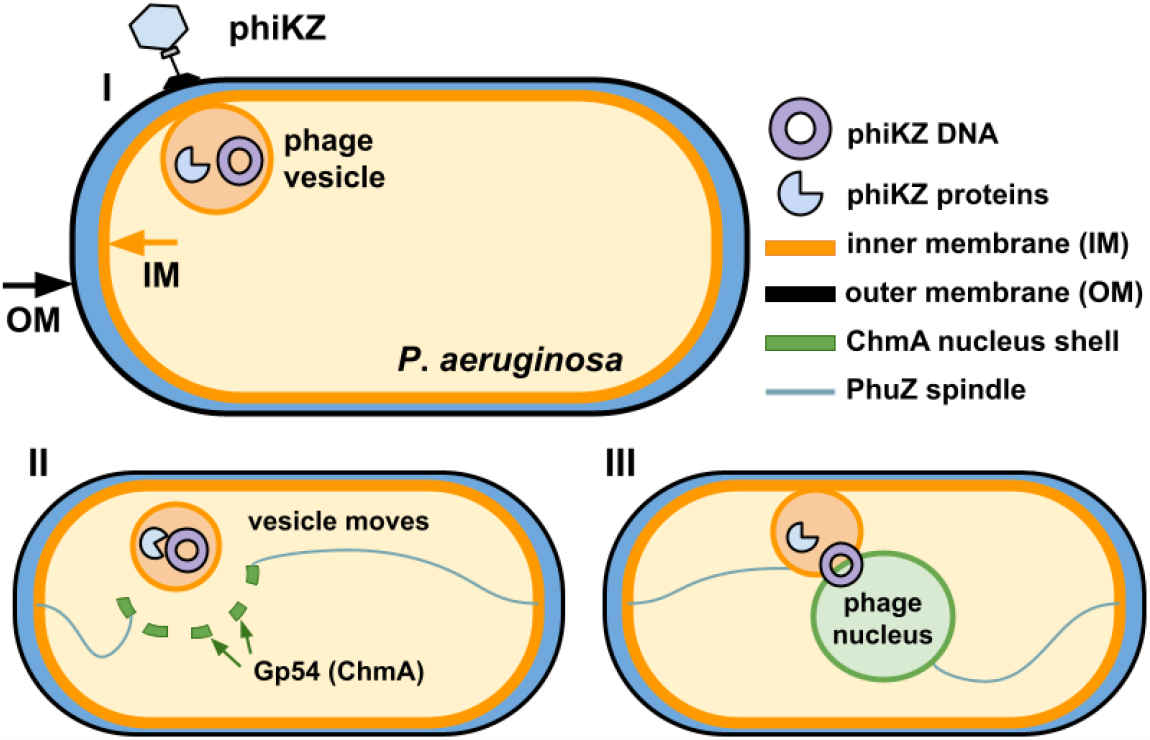
The three-stage model of phiKZ bacteriophage early infection. In the first stage (I) phage attaches to P. aeruginosa cell and injects its DNA and some virion proteins, the second stage (II) is the stage of transfer of EPI vesicle and ChmA, and the third stage is the stage of phage DNA transfer inside the phage nucleus for replication (III).

Nevertheless, there are a lot of unknown details about the functionality of the EPI vesicles. Whether the EPI vesicle provides selective protein transport from the bacterial cytoplasm inside or whether it functions only as the shield for nucleic acids is uncertain. A previous study on the *P. aeruginosa* immune system conferring resistance to phiKZ infection demonstrated that Juk proteins target the early stage of phiKZ infection and inhibit phage propagation (8). Moreover, Juk proteins were positioned with viral DNA, and the phage genome degrades (8). This may result from the interaction of the Juk proteins with the EPI vesicle membrane or penetrating the vesicle. Moreover, one of the most interesting questions is how the phage DNA moved to the phage nucleus without the total destruction of the EPI vesicle. So, the possibility of the EPI vesicle carrying out selective protein transport and details of the DNA translocation process from the EPI vesicle to the phage nucleus remains to be identified.

## Acknowledgments

TEM studies were carried out at the Shared Research Facility “Electron microscopy in life sciences” at Moscow State University (Unique set-up “Three-dimensional electron microscopy and spectroscopy”). The fluorescent microscopy experiments were carried out using the scientific equipment of the Center of Shared Usage «The analytical center of nano- and biotechnologies of SPbPU».

