## Supplementary materials for "Genomic Transfer via Membrane Vesicle: A Strategy of Giant Phage phiKZ for Early Infection"

**Supplementary Material**

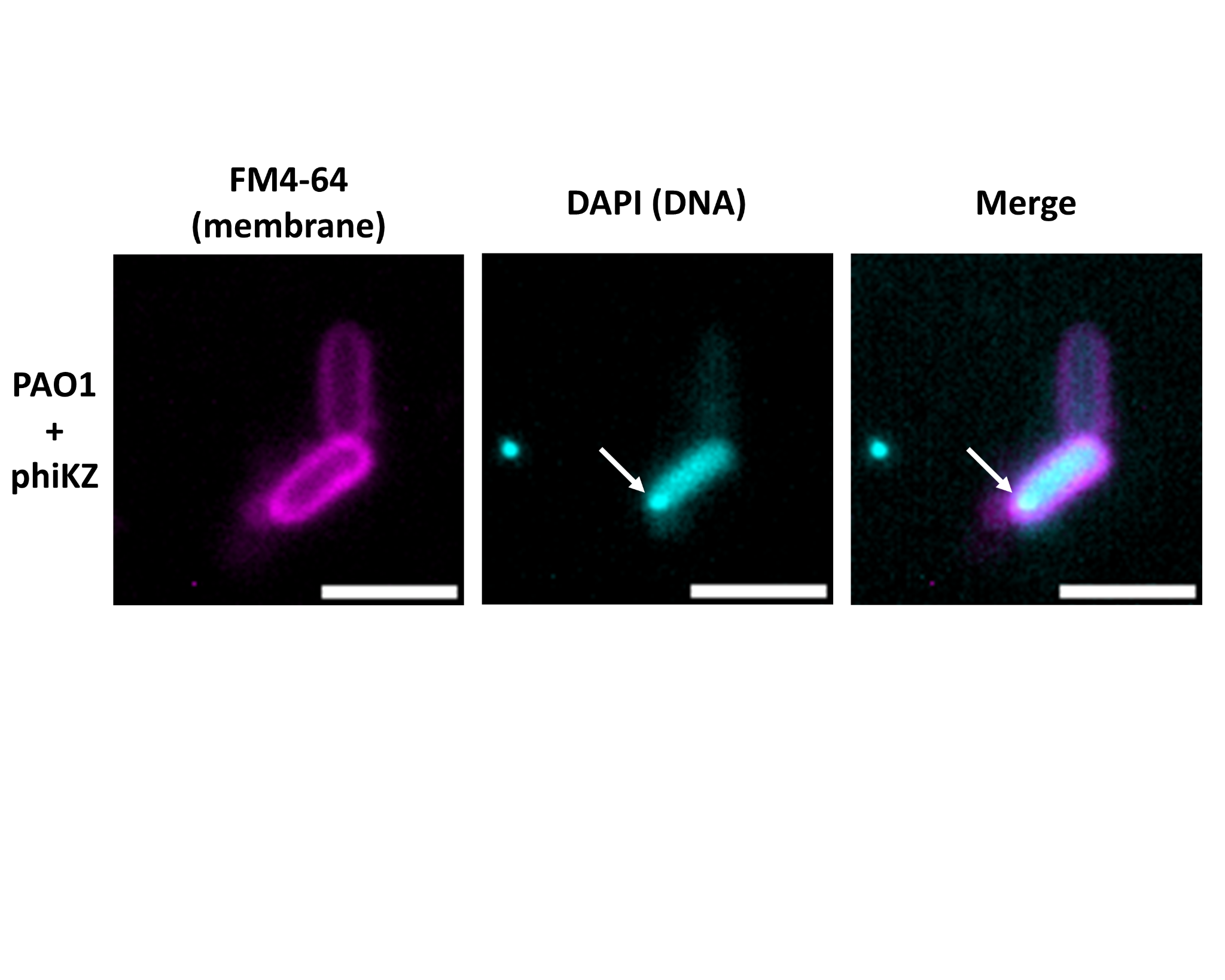

**Figure S1.** Fluorescence microscopy images of phiKZ-infected P. aeruginosa PAO1 cells stained by FM4-64 without addition of EDTA. No EPI vesicles stained by FM4-64 were observed. DAPI-stained phiKZ DNAs are indicated by white arrows. Scale bar is 2 um.

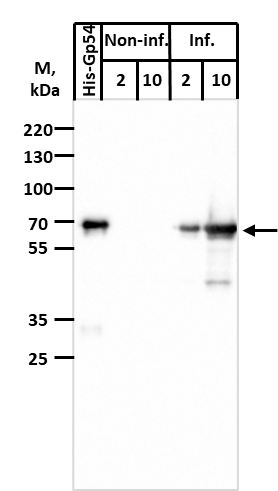

**Figure S2.** The specificity test of purified Anti-Gp54 antibodies (ABs). Purified recombinant His-Gp54 was loaded at 200 ng per lane. As a non-inf. samples were used PAO1 cells (OD600 0.65), collected before phage addition. As the "Inf." sample was used PAO1 cells after 30 min of infection by phiKZ (OD600 0.75). Samples were prepared as follows: cells from 1 ml were collected by centrifugation on 5000 g for 2 min. The sediment was resuspended in 1X Laemmli based on the ratio: 60 μl 1X Laemmli at OD600 = 0.6 per 1 ml. The numbers "2" and "10" indicate the volume in μl loaded per lane. Everywhere dilution of primary antibodies was 1:10000. Anti-Rabbit IgG (Sigma Aldrich) with the dilution of 1:10000 were used as secondary antibodies.
